# Assessment of Software Methods for Estimating Protein-Protein Relative Binding Affinities

**DOI:** 10.1101/2020.09.30.320069

**Authors:** Tawny R. Gonzalez, Kyle P. Martin, Jonathan E. Barnes, Jagdish Suresh Patel, F. Marty Ytreberg

**Author notes:** Corresponding authors, (JSP), (FMY). These authors contributed equally to this work (Tawny R. Gonzalez, Kyle P. Martin).

## Abstract

A growing number of computational tools have been developed to accurately and rapidly predict the impact of amino acid mutations on protein-protein relative binding affinities. Such tools have many applications, for example, designing new drugs and studying evolutionary mechanisms. In the search for accuracy, many of these methods employ expensive yet rigorous molecular dynamics simulations. By contrast, non-rigorous methods use less exhaustive statistical mechanics, allowing for more efficient calculations. However, it is unclear if such methods retain enough accuracy to replace rigorous methods in binding affinity calculations. This trade-off between accuracy and computational expense makes it difficult to determine the best method for a particular system or study. Here, eight non-rigorous computational methods were assessed using eight antibody-antigen and eight non-antibody-antigen complexes for their ability to accurately predict relative binding affinities (ΔΔ*G*) for 654 single mutations. In addition to assessing accuracy, we analyzed the CPU cost and performance for each method using a variety of physico-chemical structural features. This allowed us to posit scenarios in which each method may be best utilized. Most methods performed worse when applied to antibody-antigen complexes compared to non-antibody-antigen complexes. Rosetta-based JayZ and EasyE methods classified mutations as destabilizing (ΔΔ*G* < −0.5 kcal/mol) with high (83-98%) accuracy and a relatively low computational cost for non-antibody-antigen complexes. Some of the most accurate results for antibody-antigen systems came from combining molecular dynamics with FoldX with a correlation coefficient (*r*) of 0.46, but this was also the most computationally expensive method. Overall, our results suggest these methods can be used to quickly and accurately predict stabilizing versus destabilizing mutations but are less accurate at predicting actual binding affinities. This study highlights the need for continued development of reliable, accessible, and reproducible methods for predicting binding affinities in antibody-antigen proteins and provides a recipe for using current methods.

## Introduction

Protein-protein binding is an essential physiological event that governs a large number of biological processes in the cell [1]. Amino acid mutations of these proteins can introduce diversity into genomes, and disrupt or modulate protein-protein interactions by changing the underlying binding free energy (Δ*G,* i.e. binding affinity), the amount of energy required to form protein complexes [2]. The binding free energy associated with a protein-protein complex determines the stability of the complex formation and the conditions for protein-protein association. Accurate prediction of binding free energies allows us to understand how these affinities can be modified, and leads to a more comprehensive understanding of protein interactions in living organisms [3].

Experimental biophysical methods can quantitatively measure change in the protein-protein binding free energy due to a mutation (i.e. relative binding affinity, ΔΔ*G*), but these methods are typically costly, laborious, and time-consuming since all mutant proteins must be expressed and purified. Many researchers have developed and utilized computational methods to predict ΔΔ*G* values for single- or multiple-amino acid mutations (see e.g. [4–6]). Historically, the most promising in terms of accuracy are rigorous methods based on statistical mechanics that use molecular dynamics (MD) simulations and thus automatically address conformational flexibility and entropic effects [7, 8]. However, these methods are computationally expensive since they employ rigorous sampling and utilize classical mechanics [9] or quantum mechanics [10] approximations of intermolecular interactions, and require a large number of calculations per time-step. Because of the expense, rigorous methods are not well-suited to studying large sets of mutations or large proteins thus necessitating less expensive, non-rigorous methods.

Non-rigorous high-throughput methods attempt to lower the computational cost, as compared to rigorous methods, while still providing accurate ΔΔ*G* predictions. They accomplish this by including precalculated physico-chemical structural information in combination with predictive algorithms. The core mechanics that drive these methods fall under numerous classification umbrellas which have been covered by review articles [11, 12]. These review articles provide a broad overview but do not provide an unbiased, rigorous, comparative analysis outside of what the original developers provide. The developers of any given method tend to provide comparisons with other methods of the same general class to define where their method fits in the current landscape. BindProfX, for example, is available as a web server and standalone and utilizes structure-based interface profiles with pseudo counts. Upon release, it was most notably compared to FoldX (a semi-empirical trained method [13]) and DCOMPLEX (a physics-based method [14]) [15, 16]. iSEE, a statistically trained method based on 31 structure, evolution, and energy-based terms was tested against FoldX, BindProfX, and BeAtMuSiC (a machine learning-based approach [17]). Mutabind [18] and some other methods not explored in this work follow a similar testing methodology [19–21]. While these comparisons are beneficial in providing context for how a given model fits in the existing research landscape, they are not very robust, since only a narrow subset of methodologies are included. Conversely for folding stability, Kroncke et al. compared a large number of available software methods on a small dataset of transmembrane proteins providing a general overview of performance [6]. Despite the narrow dataset, this study provides a diverse, useful collection of evaluation metrics between multiple classes of methods. Our intent in this study is to provide a similar robust comparison of methods for non-rigorous binding affinity estimation.

In this work, we evaluate the ability of eight non-rigorous methods to predict relative binding affinities due to single amino acid mutations. We restrict our study to cases where both an experimental structure of the complex, and experimentally determined binding affinity values are available. To investigate the trade-off between speed and accuracy, we chose 16 protein-protein test complexes with empirical ΔΔ*G* values for observed mutations. We calculated the ΔΔ*G* values for each mutation using all eight methods and compared the results against empirical ΔΔ*G* values. The goal of this study was to determine whether software methods that use (most costly) energy functions with a wider variety of physico-chemical structural features would provide more accurate binding affinity and interface destabilization predictions compared to those that rely on a single descriptive (less costly) energy function. We have determined scenarios in which some of these methods may be better or worse than traditional computational methods in predicting ΔΔ*G* values.

## Methods

### Compilation of Experimental ΔΔG Values

To assess the performance of a range of protein-protein binding affinity prediction methods, we first assembled a dataset containing single amino acid mutations with known experimental ΔΔ*G* values. This list was assembled from Structural Kinetic and Energetic database of Mutant Protein Interaction (SKEMPI) version 2.0 [22]. While generating this list, we considered four aspects: (i) type of protein-protein complex; (ii) availability of quality 3-D structural information; (iii) range of experimental ΔΔ*G* values; and (iv) the type of mutations at differing sites on the complex. Our final dataset contained 654 mutations from 16 protein-protein complexes and their respective experimental ΔΔ*G* values. We further categorized these 16 complexes as either non-antibody-antigen (non-Ab) or antibody-antigen (Ab). Table 1 shows the complexes in our dataset with their respective non-Ab and Ab categories and the number of mutations associated with each complex. The dataset contains a total of 401 non-Ab mutations and 253 Ab mutations.

**Table 1.** Dataset used in our study containing 16 protein complexes. For both non-Ab (left) and Ab (right) categories, columns show PDB IDs, total number of residues in a complex, and number of experimental mutants per complex.

| Non-Ab |  |  | Ab |  |  |
| --- | --- | --- | --- | --- | --- |
| PDB ID | # Mutations | # Residues | PDB ID | # Mutations | # Residues |
| 1a4y [23] | 32 | 583 | 1bj1[24] | 10 | 547 |
| 1brs [25] | 30 | 199 | 1jrh [26] | 42 | 540 |
| 1cbw [27] | 31 | 299 | 1mlc [28] | 11 | 561 |
| 1iar [29] | 36 | 336 | 1vfb [30] | 48 | 352 |
| 1jtg [31] | 37 | 428 | 1yy9 [32] | 16 | 1058 |
| 1lfd [33] | 19 | 254 | 2jel [34] | 43 | 520 |
| 1ppf [35] | 190 | 274 | 3hfm [36] | 71 | 558 |
| 2wpt [37] | 26 | 220 | 4i77 [38] | 12 | 549 |

### Selection of Protein-Protein Binding Affinity Methods

Binding affinity prediction methods were chosen to have both a distinct approach to binding affinity calculation that utilized 3-D structural information and had functional standalone software in September 2020, available either online or upon request to the author. Table 2 summarizes the methods selected in this study, their approaches, and their type of scoring functions. For simplicity, we categorized scoring functions (mathematical functions to calculate ΔΔ*G* values) as semi-empirical, statistical, or physics-based. Semi-empirical methods replace as many calculations as possible with pre-calculated data and are trained using existing crystal structures and known binding affinity measurements for mutations [39]. Statistical methods use pre-calculated data and consider changes in coarse structural features such as the change in overall volume [40]. Physics-based methods use molecular mechanics based-energy functions to estimate enthalpic binding contributions [14]. In general, statistical or semi-empirical scoring functions involve a training step where existing datasets are leveraged to determine the weight of input parameters. MD, JayZ, and EasyE were not developed by training the methods against experimental data designed to improve predictive power while all other methods utilized this step.

**Table 2.** Methods used for comparison in study with a short summary of their approach and scoring function. Columns (left to right) indicate the method, a brief description of the method, the type of scoring function used, and runtimes. Runtimes are the amount of CPU hours for estimating the ΔΔ*G* for a representative protein complex for Ab (1yy9, 1058 residues) and Non-Ab (1ppf, 274 residues) categories. Although 1yy9 is roughly four times bigger than 1ppf, the total runtime may or may not be affected depending on the method used.

| Name | Brief Description | Scoring Function | Runtime (CPU hours) |
| --- | --- | --- | --- |
| BindProfX [15, 16] | Interface profile score based on conservation of homologous interfaces | Semi-Empirical | 1ppf = 0.57 CPUh<br>1yy9 = 0.73 CPUh |
| BindProfX(BPX)+FoldX v4 [15, 16] | Profile score weighted and combined with FoldX energy potential | Semi-Empirical | 1ppf = 0.62 CPUh<br>1yy9 = 0.71 CPUh |
| iSEE [41] | Random forest model using structural, evolutionary, and energy-based features | Statistical | 1ppf < 0.01 CPUh<br>1yy9 < 0.01 CPUh |
| DCOMPLEX v2 [14] | Structural ideal-gas reference state potential | Physics-Based | 1ppf = 0.013 CPUh<br>1yy9 = 0.001 CPUh |
| EasyE v1.0 [40, 42] | GMEC-based method utilizing the Rosetta [43, 44] energy function | Statistical | 1ppf = 0.48 CPUh<br>1yy9 = 0.09 CPUh |
| JayZ v1.0 [40, 42] | Partition-function method utilizing Rosetta energy function | Statistical | 1ppf = 0.14 CPUh<br>1yy9 = 0.21 CPUh |
| FoldX v4 [13, 39] | Empirical energy score based on various energy parameters (e.g. van der Waals, solvation, electrostatics, hydrogen bonding) | Semi-Empirical | 1ppf = 0.42 CPUh<br>1yy9 = 0.16 CPUh |
| MD+FoldX v4 [45-47] | Molecular dynamics used to explore conformation space and generate snapshots; FoldX score calculated for each snapshot and averaged | Semi-Empirical | 1ppf = 941 CPUh<br>1yy9 = 4093 CPUh |

### Calculation and Comparison of Computational Speed

The methods in Table 2 were used to predict ΔΔ*G* values for each mutation on our experimental list shown in Table 1. Detailed protocols for predicting ΔΔ*G* values using each selected method are provided in the Supplemental Information (see S1 File). Runtimes were determined by using a representative protein complex from each category: 1ppf, a non-Ab complex with 274 total amino acids, and 1yy9, an Ab complex with 1058 total amino acids (see Table 2). These runtimes are estimates provided to give a point of comparison between the speeds of different methods. Specific runtimes will be determined by hardware specifications, method parameters, the number of mutations being computed, and overall protein size. For MD+FoldX, computational runtime was the length of time of the MD simulation plus the FoldX runtime for a single mutation. Reporting runtime in this fashion highlights the large CPUh requirement needed in order to add the sampling of MD into FoldX calculations. We note that, in contrast to the other methods tested here, the MD simulations that must be performed for MD+FoldX can be completed very quickly on modern GPUs, significantly offsetting the high initial cost of the MD+FoldX method. For all other methods, the algorithms rely either on various pre-calculated data or limited conformational sampling to calculate ΔΔ*G* values rapidly.

### Comparing Experimental and Predicted ΔΔ*G* Values

To carry out statistical analysis of our results we built an in-house Python script (see S2 File) that uses a combination of libraries including matplotlib, numpy, pandas, statistics, scipy, and sklearn. Using this script, we compared predicted values to experimental ΔΔ*G* values for each method.

To evaluate the predictive ability of each method tested, we compared the following correlation coefficients using our script: concordance (*ρ_c_*), Pearson (*r*), Kendall (τ), and Spearman (*ρ*) (see Table 3). We distinguish between methods that were trained to predict ΔΔ*G* values from methods that compute metrics that are expected to linearly correlate with ΔΔ*G* values. This distinction is important since for optimal performance we expect a regression line that passes through the coordinate origin and has a slope of 1, leading to a correlation coefficient equal to 1.

**Table 3.** Statistical measures used to test the performance of each method in predicting ΔΔ*G* values.

| Correlation | Brief Description | Type |
| --- | --- | --- |
| Concordance | The concordance correlation coefficient ($\rho_c$) measures the degree to which the predicted $\Delta\Delta G$ value equals the actual experimental value (0 indicates no agreement and 1 perfect agreement). | Linear |
| Pearson | The Pearson correlation coefficient ($r$) measures the degree to which a uniform linear transformation of the predicted $\Delta\Delta G$ values (i.e., a shift and scale change) would yield the actual experimental values (0 indicates no agreement after transformation, 1 perfect agreement, and $-1$ perfect inverse agreement). | Linear |
| Kendall and Spearman | The rank correlation coefficient measures the degree to which the rank ordering of the predicted $\Delta\Delta G$ values matches the rank ordering of the actual experimental values (0 indicates no agreement after transformation, 1 perfect agreement, and $-1$ perfect inverse agreement). In a normal case, the Kendall correlation ($\tau$) is considered more robust than the Spearman correlation ($\rho$) because of a smaller gross error sensitivity and more efficient due to a smaller asymptotic variance [48]. | Rank |
| AUC and ROC | The receiver operating characteristic (ROC) curve tests several cutoff values for binning mutations as neutral or destabilizing between the most negative calculated $\Delta\Delta G$ value and the most positive calculated $\Delta\Delta G$ value, with true positive rates (sensitivity) calculated at each point. As the true positive rate is calculated, the classifier is moved to less extreme values; this yields the ROC curve. The area under curve (AUC) is a summary statistic that approximates how well the predictor actually discriminates between the two classifications. | N/A |

To compare the discriminating power of the methods, we generated receiver operating characteristic (ROC) curves (see Table 3). These curves quantify the ability of a method to correctly classify point mutations as destabilizing (ΔΔ*G* < −0.5 kcal/mol) or neutral/stabilizing (ΔΔ*G* > −0.5 kcal/mol). ROC curves that are skewed toward a higher true positive rate (sensitivity) classify mutations more accurately, as quantified by area under curve (AUC, ranging between 1.0 and 0.5 for perfect and chance classification, respectively).

We also used our script to parse the results on the basis of several physico-chemical and structural features to allow us to evaluate the methods based on these characteristics: wild type amino acid type, mutant amino acid type, protein-protein interacting versus antibody-antigen, secondary structure classification of the mutation [49, 50], coordination number [51], Sneath index [52], mostly α-helical proteins versus mostly β-sheet proteins versus a mix of both α-helical and β-sheet proteins, percent exposure, location of the mutation, change in charge, change in polarity, change in volume, and whether or not the mutation location is predicted as an active or passive residue [53–55]. The script uses data from S3 File as an input and outputs scatter plots, correlation plots, receiver operating characteristic (ROC) curves, and box plots to visualize the data, as well as correlations and standard deviations for each method. All plots in this manuscript were generated using this script.

## Results

The purpose of our study was to assess the ability of eight different relative binding affinity calculation methods (see Table 2) to estimate ΔΔ*G* values. We selected 16 different protein complexes (eight Ab, eight non-Ab, see Table 1) with a total of 654 single amino acid mutations. Each method was then used to estimate ΔΔ*G* values of 654 mutations and a variety of statistical measures were employed to assess their predictive ability. We also examined the computational speed of each method in the context of accuracy to determine its efficiency.

### Non-Antibody-Antigen (non-Ab) Results

Our dataset of eight non-Ab test protein complexes contains 401 total mutations and are mainly classified as protein-protein systems formed by inhibitors and receptors that range from 199 to 583 residues in size. The distribution and our classification of experimental ΔΔ*G* values for all non-Ab test complexes is as follows: 13% of point mutations resulted in ΔΔ*G* values less than −0.5 kcal/mol (classified as destabilizing); 31% between −0.5 and 0.5 kcal/mol (neutral); and 56% greater than 0.5 kcal/mol (stabilizing).

Figures 1 (blue data points and values) and 2 show various performance metrics for each method to assess their ability to predict the non-Ab ΔΔ*G* values. Overall, EasyE has the highest correlation coefficient, *r* = 0.62, and iSEE has the lowest, *r* = 0.17 (see Figures 1 and 2). JayZ and EasyE, both of which utilize Rosetta’s conformational sampling algorithms, consistently have the best *r* values for non-Ab mutations.

**Figure 1.**
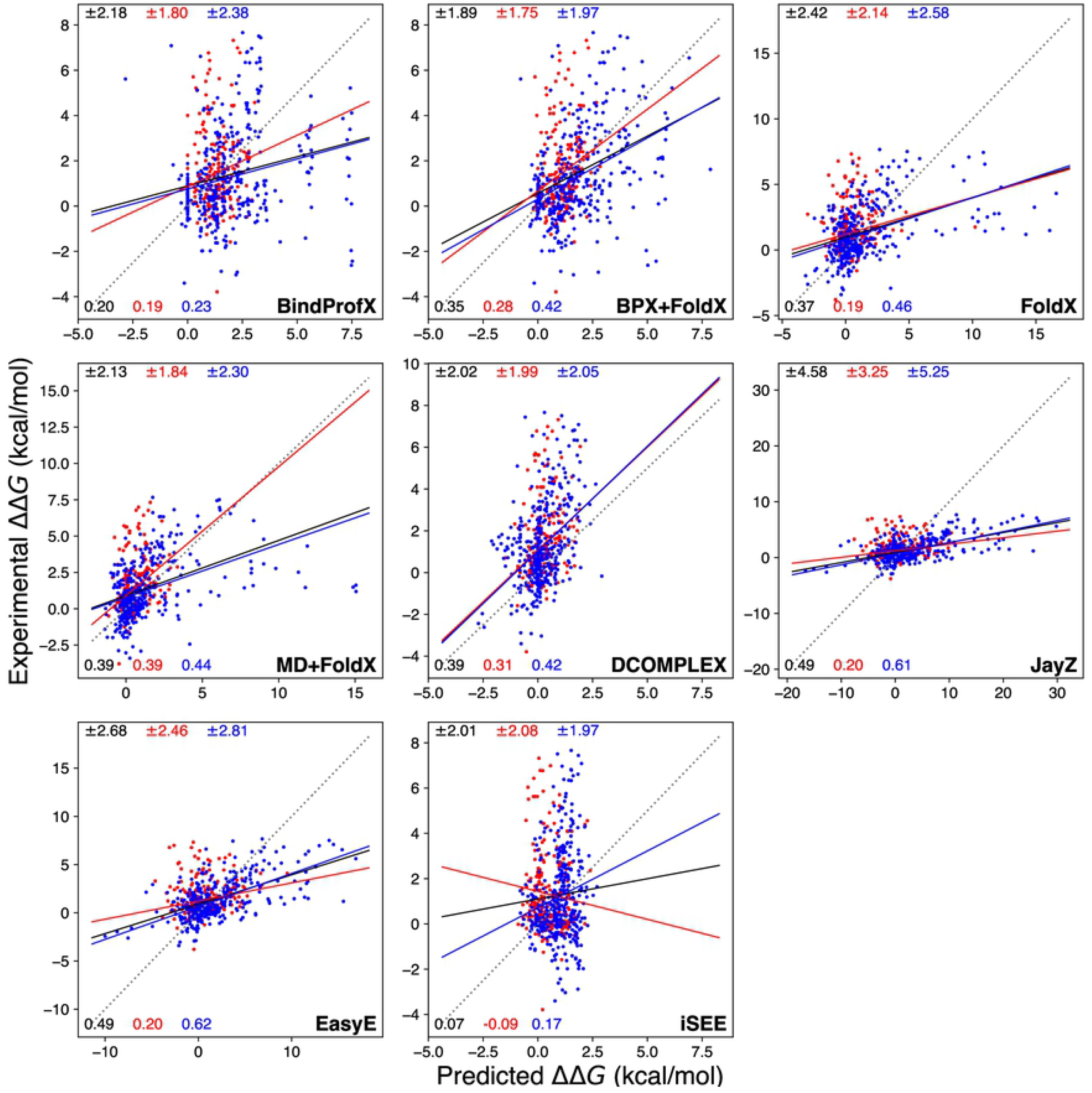
Calculated ΔΔ*G* values (x-axis) compared to experimental ΔΔ*G* values (y-axis) for each method tested in this study. Black, red, and blue lines are simple linear regressions from which *r* are derived. The red points are a scatter for Ab complexes and the blue points are for non-Ab complexes. The dashed line is the *y = x* line measuring perfect agreement between predicted and experimental ΔΔ*G* values. The solid black, red, and blue lines indicate a linear relationship between calculated and experimental observations for all data points, Ab complexes, and non-Ab complexes respectively. The top values in black, red, and blue match the root-mean-square error and the bottom values indicate *r* for all values, Ab values, and non-Ab values respectively.

**Figure 2.**
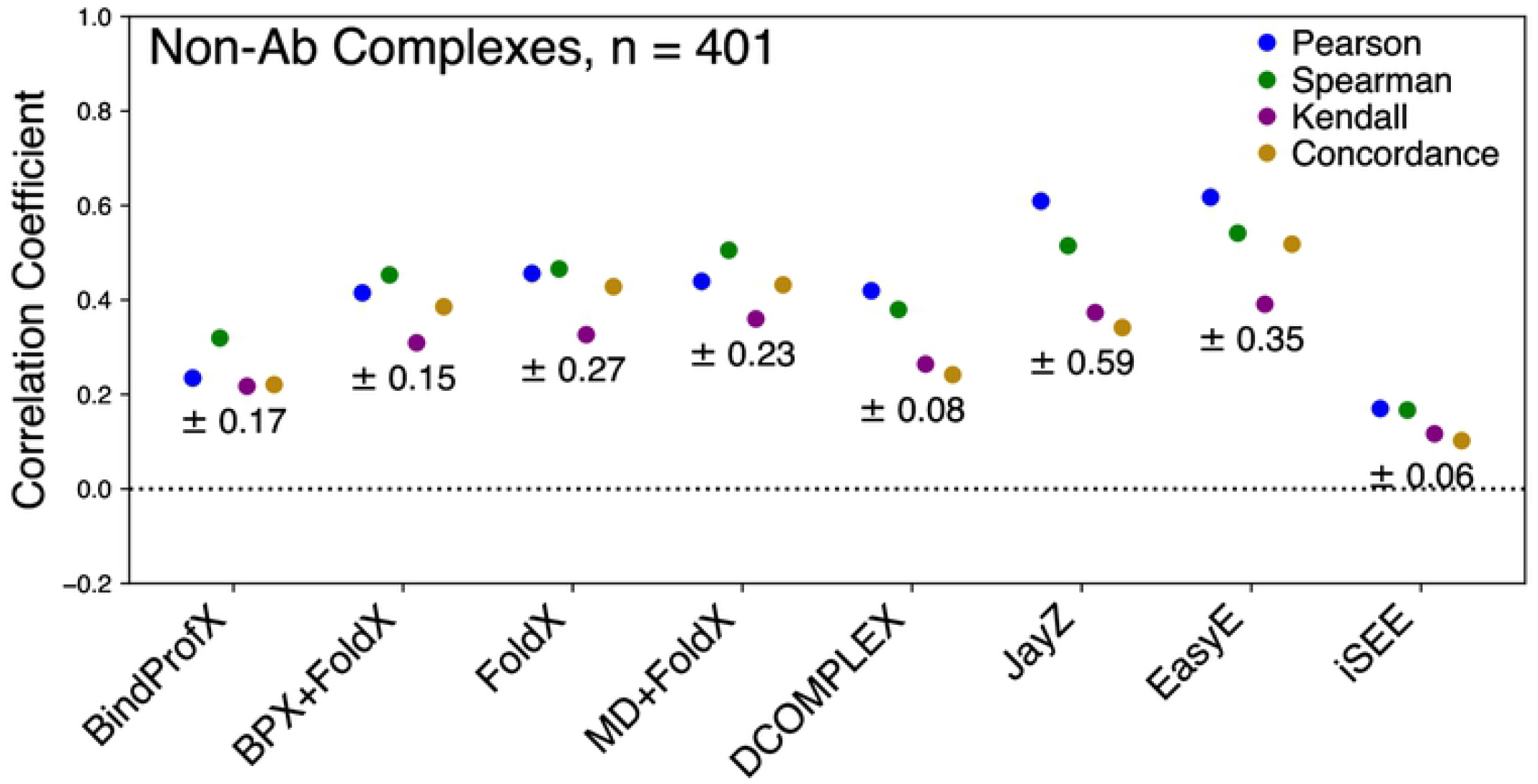
Performance of each method for non-Ab complexes (401 total mutations) in predicting true ΔΔ*G* values (*ρ_c_*), linearly correlated ΔΔ*G* values (*r*), and rank order (*ρ* and τ). The error for each method is reported under the correlation points.

Figure 3 shows the ROC plot for all the tested methods. These ROC plots highlight how well a method can discriminate between stabilizing and destabilizing mutations. JayZ (0.84), EasyE (0.83), DCOMPLEX (0.82), FoldX (0.79), and MD+FoldX (0.76) have the highest AUC. Combined with the results from Figures 1 and 2, for the systems studied here, JayZ and EasyE methods are the best overall performers in terms of accuracy, discriminating stabilizing mutations from destabilizing, and ranking mutations based on their experimental ΔΔ*G* values.

**Figure 3.**
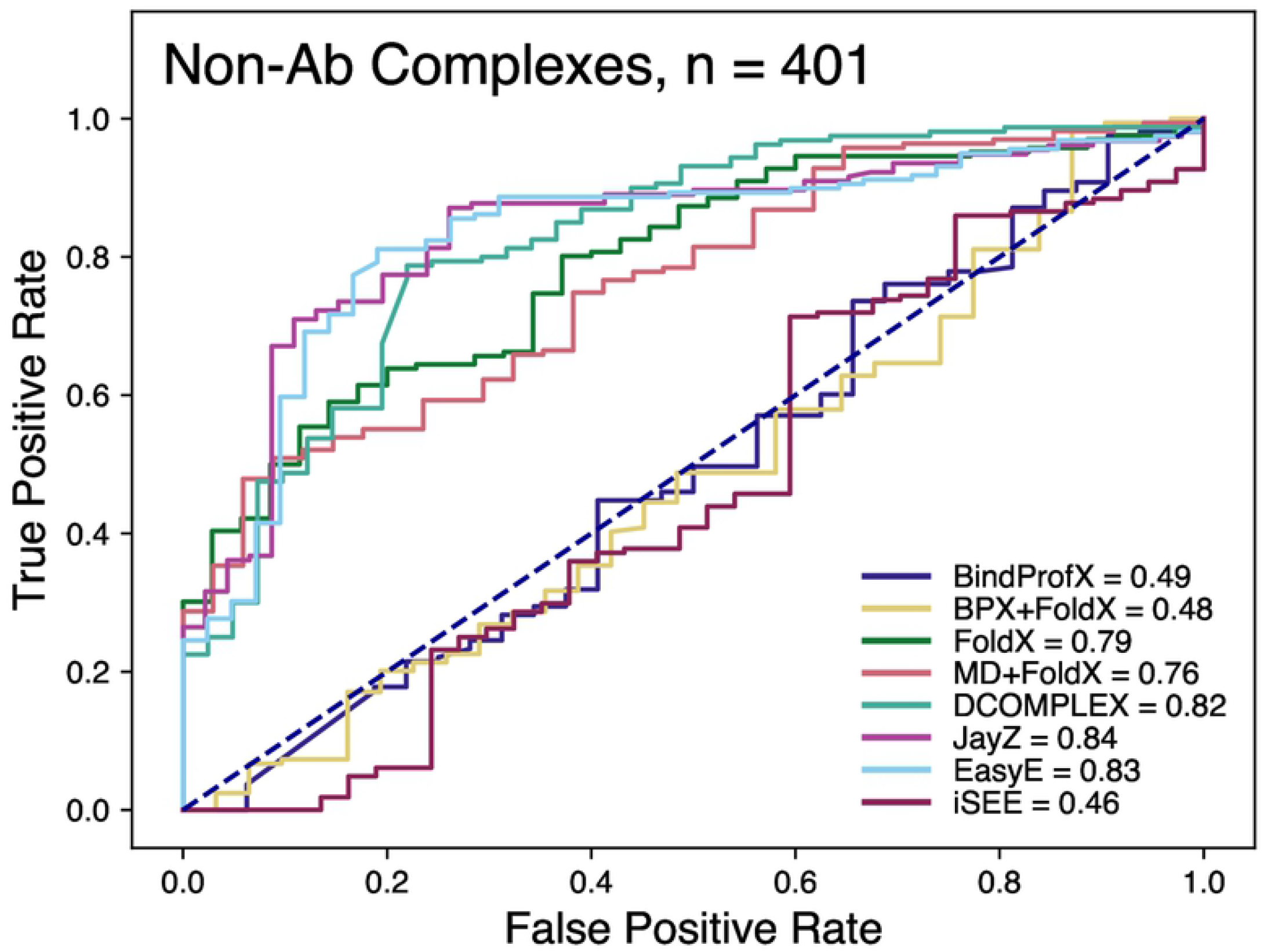
Receiver operating characteristic (ROC) curves for non-Ab complexes of the classification of variants as stabilizing (ΔΔ*G* < −0.5 kcal/mol) or destabilizing (ΔΔ*G* > 0.5 kcal/mol). The values in the legend represent the area-under-curve (AUC). The higher the value, the better method is at discriminating between destabilizing and destabilizing mutations.

Table 2 reports CPUh required (i.e. runtimes) for each method to calculate ΔΔ*G* for the entire list of mutations for a representative non-Ab protein complex. BindProfX, BindProfX(BPX)+FoldX, JayZ, and EasyE allow users to specify a list of mutations that the method is then able to calculate in one setting. This list can be optimized based on the available hardware to achieve efficiency. iSEE requires significant preparatory work (see File S1) prior to calculation, but once completed, it calculates the ΔΔ*G* values for the entire list of mutations nearly instantly. DCOMPLEX is not as flexible out of the box but can handle large numbers of mutations through an automated script. For MD+FoldX, 1yy9 (roughly four times larger than 1ppf) requires considerably more CPUh to calculate. All other methods calculate 1yy9 in a shorter time frame than 1ppf. This may seem counterintuitive. However, MD must statistically sample the conformational energy of the entire complex, while all other methods use algorithms that are likely impacted more by the number of residues involved in the interaction rather than the protein size. Overall, DCOMPLEX has a much faster runtime compared to other methods, and if the goal is to determine stabilizing and destabilizing non-Ab mutations, it offers similar discriminating power to JayZ and EasyE, at a fraction of the computational cost. JayZ estimates ΔΔ*G* value of one mutation in ~2.7 s, EasyE in ~9.1 s, but DCOMPLEX requires just ~0.25 s. Overall, EasyE appears to be the best option for balancing accuracy and speed and DCOMPLEX is recommended for discriminating between stability and destabilizing mutations.

A method might not be a good overall performer in predicting ΔΔ*G* values but could still perform well for mutations with certain physico-chemical and structural features. Therefore, we calculated various statistical measures to assess each method on unique subsets of mutations (see Table 4 and SI Figs S1-4). This table shows eight different data subsets with two *r* per method. EasyE has the highest *r* for non-Ab for five out of eight subsets (wild type non-gly or non-pro, alpha helix, beta sheet, surface exposure, and large volume changes). Where this method did not have the highest *r*, it had either the second or third highest *r*. JayZ mirrors the performance of EasyE in all the same categories and performs better than Easy in the neutral charge subset. These results further highlight the versatility of EasyE’s and JayZ’s performance in estimating the effects of non-Ab mutations compared to the other methods tested in this study. All methods apart from iSEE and BindProfX perform surprisingly well in the WT Gly or Pro subset. iSEE’s performance in this subset, while still mediocre compared to the other tested methods, is substantially better than in all other subsets.

**Table 4.** All methods *r* with respect to certain subsets. “WT Gly or Pro” are wild type amino acids that are either glycine or proline. “WT Non-Gly or Non-Pro” are wild type amino acids that are neither glycine nor proline. “Alpha Helix” are mutations that occur in a helix structure. “Beta Sheet” are mutations that occur in a beta structure. “Surface Exposure” are mutations that occur in an amino acid that have relative solvent accessibility values between 0 and 10%. “Neutral Charge” is a neutrally charged wild type amino acid mutating to a neutrally charged mutant amino acid. “Hydrophobic to Polar” is a hydrophobic or polar wild type amino acid mutating to a polar or hydrophobic mutant amino acid, respectively. “Larger Vol Changes” is a mutant amino acid that is greater than 40% larger than the wild type amino acid. Values that are bolded are the highest *r* for each method and protein type. Values that are red or blue are the highest *r* for each subset, blue for non-Ab and red for Ab.

| Method | WT Gly or Pro | WT Non-Gly or Non-Pro | Alpha Helix | Beta Sheet | Surface Exposure | Neutral Charge | Hydrophobic to Polar | Large Vol Changes |
| --- | --- | --- | --- | --- | --- | --- | --- | --- |
| BindProfX | Non-Ab: 0.11<br>Ab: -0.03 | Non-Ab: 0.33<br>Ab: 0.23 | Non-Ab: 0.29<br>Ab: 0.16 | Non-Ab: 0.29<br>Ab: **0.52** | Non-Ab: 0.22<br>Ab: 0.09 | Non-Ab: **0.37**<br>Ab: 0.28 | Non-Ab: 0.33<br>Ab: 0.17 | Non-Ab: 0.13<br>Ab: 0.42 |
| BPX+FoldX | Non-Ab: **0.81**<br>Ab: 0.09 | Non-Ab: 0.45<br>Ab: 0.34 | Non-Ab: 0.43<br>Ab: 0.39 | Non-Ab: 0.43<br>Ab: **0.54** | Non-Ab: 0.32<br>Ab: 0.21 | Non-Ab: 0.52<br>Ab: 0.41 | Non-Ab: 0.41<br>Ab: 0.26 | Non-Ab: 0.71<br>Ab: **0.50** |
| FoldX | Non-Ab: **0.85**<br>Ab: -0.11 | Non-Ab: 0.45<br>Ab: 0.25 | Non-Ab: 0.39<br>Ab: 0.25 | Non-Ab: 0.39<br>Ab: 0.31 | Non-Ab: 0.50<br>Ab: 0.26 | Non-Ab: 0.42<br>Ab: **0.41** | Non-Ab: 0.41<br>Ab: 0.11 | Non-Ab: 0.63<br>Ab: -0.32 |
| MD+FoldX | Non-Ab: **0.83**<br>Ab: **0.71** | Non-Ab: 0.49<br>Ab: **0.42** | Non-Ab: 0.44<br>Ab: **0.54** | Non-Ab: 0.44<br>Ab: 0.49 | Non-Ab: 0.47<br>Ab: 0.35 | Non-Ab: 0.46<br>Ab: 0.46 | Non-Ab: **0.46**<br>Ab: **0.31** | Non-Ab: 0.71<br>Ab: 0.35 |
| DCOMPLEX | Non-Ab: **0.65**<br>Ab: **0.89** | Non-Ab: 0.34<br>Ab: 0.37 | Non-Ab: 0.33<br>Ab: 0.31 | Non-Ab: 0.33<br>Ab: 0.30 | Non-Ab: 0.52<br>Ab: 0.27 | Non-Ab: 0.36<br>Ab: **0.56** | Non-Ab: 0.38<br>Ab: 0.16 | Non-Ab: 0.62<br>Ab: 0.28 |
| JayZ | Non-Ab: 0.80<br>Ab: **0.54** | Non-Ab: 0.49<br>Ab: 0.24 | Non-Ab: 0.44<br>Ab: -0.06 | Non-Ab: 0.44<br>Ab: 0.16 | Non-Ab: 0.59<br>Ab: **0.36** | Non-Ab: **0.62**<br>Ab: 0.26 | Non-Ab: 0.41<br>Ab: 0.01 | Non-Ab: **0.83**<br>Ab: 0.19 |
| EasyE | Non-Ab: 0.80<br>Ab: 0.29 | Non-Ab: **0.51**<br>Ab: 0.22 | Non-Ab: **0.51**<br>Ab: 0.06 | Non-Ab: **0.51**<br>Ab: 0.03 | Non-Ab: **0.60**<br>Ab: **0.35** | Non-Ab: 0.61<br>Ab: 0.23 | Non-Ab: 0.45<br>Ab: 0.02 | Non-Ab: **0.84**<br>Ab: 0.18 |
| iSEE | Non-Ab: **0.43**<br>Ab: -0.43 | Non-Ab: 0.28<br>Ab: -0.16 | Non-Ab: 0.05<br>Ab: -0.04 | Non-Ab: 0.05<br>Ab: -0.24 | Non-Ab: 0.15<br>Ab: **0.11** | Non-Ab: 0.15<br>Ab: -0.11 | Non-Ab: 0.14<br>Ab: -0.02 | Non-Ab: 0.24<br>Ab: -0.44 |

### Antibody-Antigen (Ab) Results

Our dataset of eight Ab test protein complexes contains 253 mutations and the proteins range in size from 352 to 1058 residues. The distribution and our classification of experimental ΔΔ*G* values for all Ab test complexes is as follows: 5% of point mutations resulted in ΔΔ*G* values less than −0.5 kcal/mol (classified as destabilizing); 40% between −0.5 and 0.5 kcal/mol (neutral); and 55% greater than 0.5 kcal/mol (stabilizing).

Figures 1 (data points and values in red), 4, and 5 show the performance of each method in predicting the ΔΔ*G* values of Ab mutations. Overall, the highest correlation is for MD+FoldX with *r* = 0.39 and the lowest is iSEE with *r* = −0.09 (see Figures 1 and 4). An interesting trend is that the methods with the highest *r* values for non-Ab complexes do not have the highest *r* for Ab complexes.

**Figure 4.**
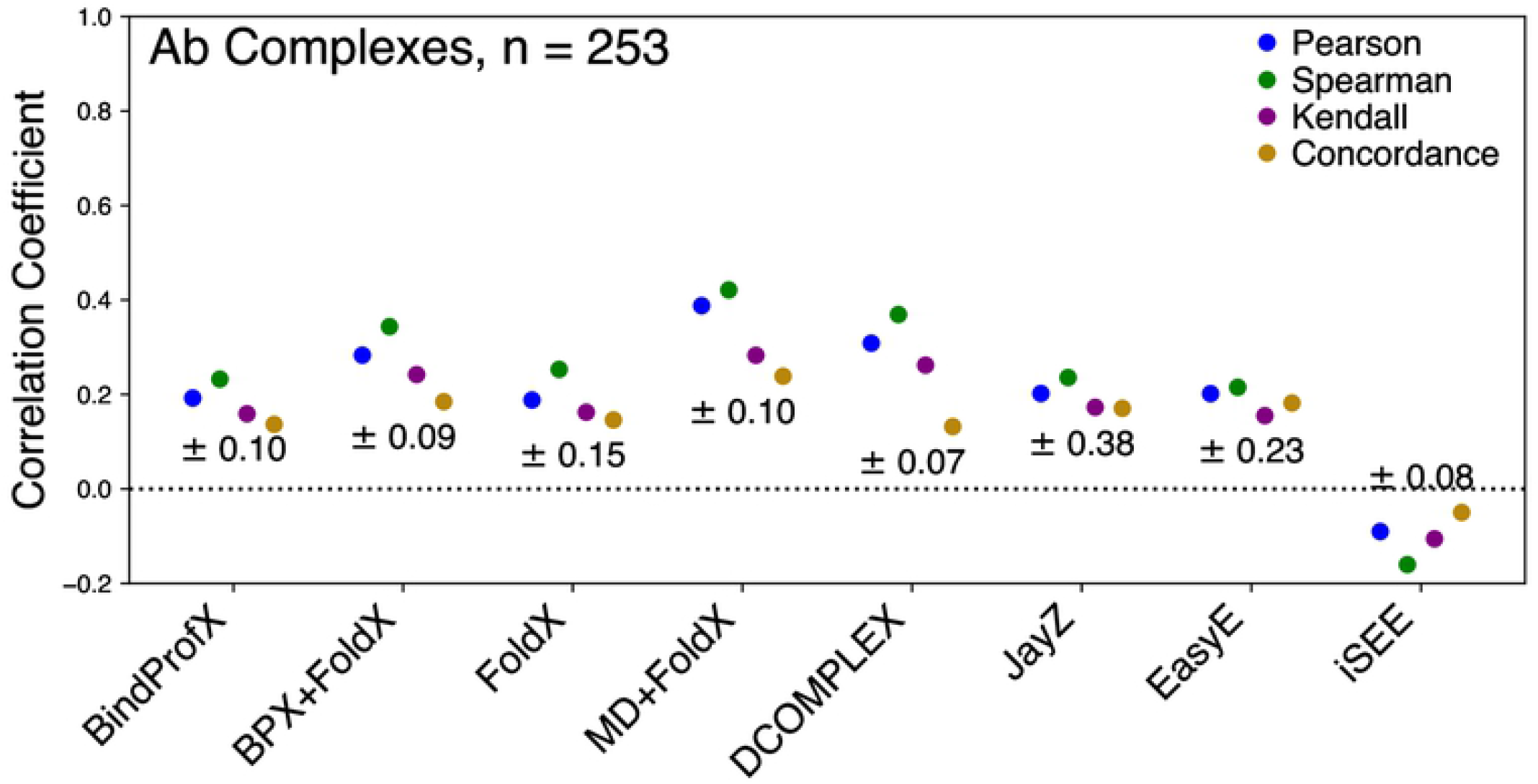
Performance of each evaluated method for Ab complexes (253 total mutations) in predicting true ΔΔ*G* values (*ρ_c_*), linearly correlated ΔΔ*G* values (*r),* and rank order (*ρ* and τ). The error for each method is reported under the correlation points.

Figure 5 shows the ROC plot for all the tested Ab methods. These ROC plots highlight how well a method is actually able to discriminate between stabilizing and destabilizing mutations. Compared to non-Ab complexes, all methods performed better for antibody-antigen complexes except for FoldX and DCOMPLEX which were marginally worse. JayZ (0.97), EasyE (0.98), FoldX (0.85), and MD+FoldX (0.82) had the highest AUC values. Combined with the results from Figures 1 and 4, at least for the systems studied here, it appears that the MD+FoldX method is the best overall performer in terms of accuracy, discriminating stabilizing mutations from destabilizing, and ranking mutations based on their experimental ΔΔ*G* values.

**Figure 5.**
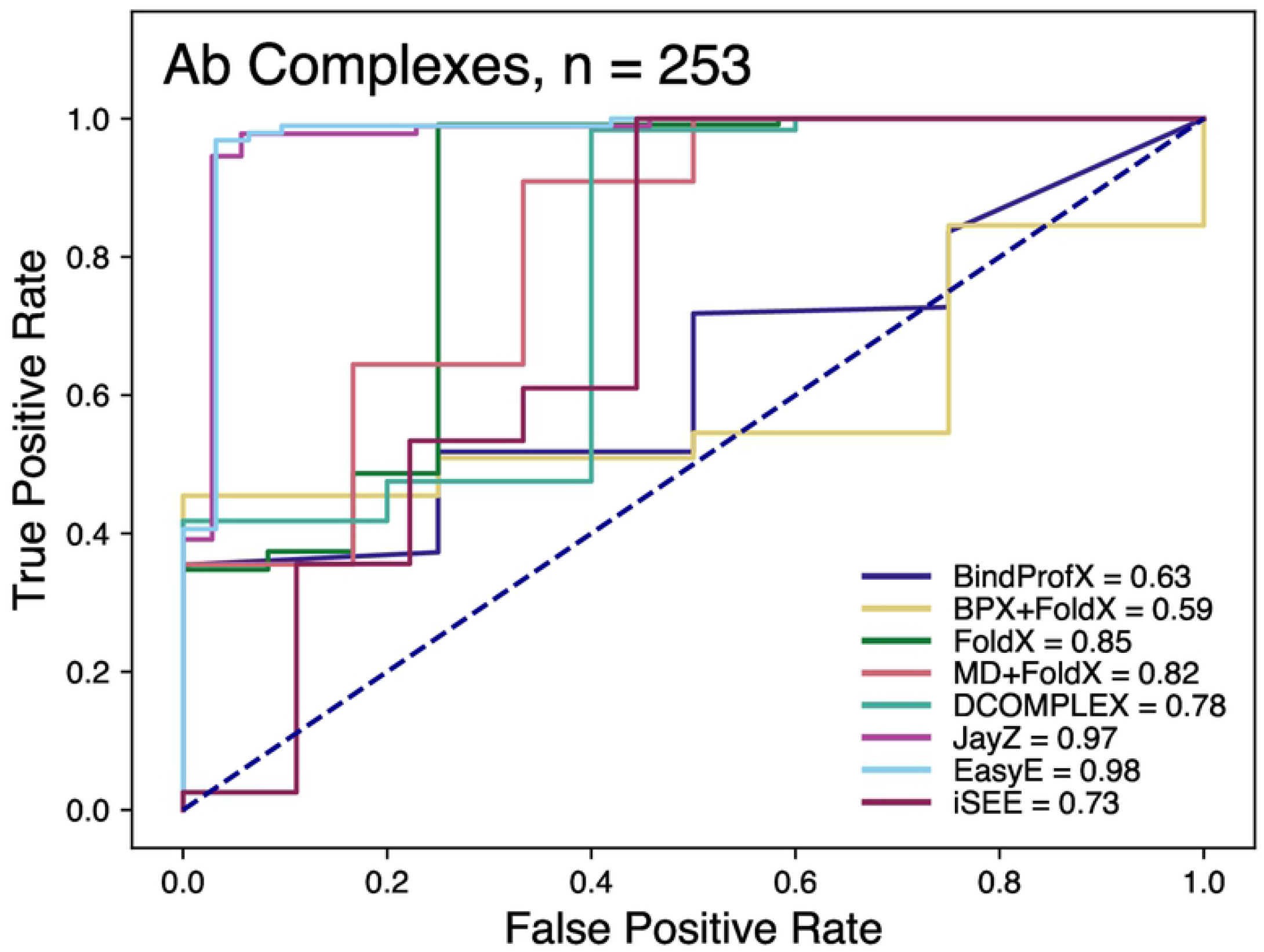
Receiver operating characteristic curves of the classification of variants that are more destabilized or less destabilized than 0.5 kcal/mol. The values in the legend represent the area-under-curve (AUC). The higher the value, the better the prediction capability of the method.

Compared to other methods, EasyE has a much faster runtime and is recommended if the goal is to discriminate between stabilizing and destabilizing (ΔΔ*G* for one mutation takes ~21 s, see Table 2). By comparison, MD+FoldX cost ~941 CPUh for one mutation of 1yy9. DCOMPLEX provides a slightly lower *r* (0.31) and computational cost (~0.35 s) for one mutation of 1yy9. Overall, MD+FoldX appears to be the best option for accuracy and EasyE or JayZ are the best options for discriminating between destabilizing and stabilizing mutations.

Table 4 summarizes the ability of each method to predict ΔΔ*G* values for subsets of Ab mutations. Most methods had mediocre *r* values less than 0.60. The exceptions to this are MD+FoldX and DCOMPLEX within the WT Gly or Pro subset with *r* = 0.71 and 0.89, respectively. MD+FoldX has the highest *r* for Ab complexes for three of the eight subsets (WT nonGly or nonPro, alpha helix, and hydrophobic to polar). BPX+FoldX has the highest *r* in two of the eight subsets (beta sheet and large volume changes). For the beta sheet subset, BindProfX had the second highest *r*. DCOMPLEX had the highest *r* for two different subsets (WT Gly or Pro and neutral charge). In the surface exposure subset, JayZ and EasyE both had nearly identical *r* (0.36 and 0.35 respectively), the highest for this subset, but substantially worse than they did for non-Ab complexes.

## Discussion

We assessed the performance of eight distinct protein-protein binding affinity calculation methods that use 3-D structural information. To test the performance of these methods, we selected 16 different protein complexes (see Table 1) with a total of 654 single amino acid mutations: eight antigen-antibody complexes (Ab, 253 mutations) and eight non-antigen-antibody (Non-Ab, 401 mutations) complexes. Each method was used to estimate ΔΔ*G* values of the 654 mutations, a variety of statistical measures, CPU cost, and physico-chemical structural features to assess the performance. Our results suggest each method has both strengths and weaknesses depending on the properties of the protein system. Most methods did not perform well when applied to mutations in Ab complexes compared to non-Ab complexes. Rosetta-based JayZ and EasyE were able to classify mutations as destabilizing (ΔΔ*G* < −0.5 kcal/mol) with high (83-98%) accuracy at relatively low computational cost. Some of the best results for Ab systems came from combining MD simulations with FoldX with a *r* coefficient of 0.39, but at the highest computational cost of all the tested methods.

Figure 1 summarizes the performance of each method in terms of its ability to estimate ΔΔ*G* values for all (non-Ab + Ab) single mutations. None of the test methods show a very high *r* between experimental and predicted ΔΔ*G* values. Two of the best performing methods, JayZ and EasyE, both have an *r* of 0.49 for all mutations, with a higher *r* of 0.61 and 0.62 respectively for non-Ab complexes. These results agree with published results from the authors of JayZ and EasyE. Our results agree moderately with published results from iSEE (they obtained *r* = 0.25, we obtained *r* = 0.17) and BindProfX (they used a much larger dataset). Published results for DCOMPLEX show a very good correlation of *r* = 0.87; much larger than what we obtained here. This difference is very likely due to the dataset size and compilation; DCOMPLEX was originally tested against 69 experimental data points, compared to the 654 values used here. MD+FoldX has an *r* of 0.39 for Ab complexes and appears to perform well for larger systems, which could indicate the importance of conformational sampling for antibody-antigen systems. Other methods used in this study have little to no conformational sampling which could explain their poor performance on Ab complexes. By contrast, these same methods perform well for non-Ab complexes, suggesting that conformational sampling is not the limiting factor to achieve accurate results for these protein complexes. For example, FoldX has a trained scoring function derived using a dataset of mostly non-Ab complexes and performs poorly for Ab complexes when using a single structure (see Table 2). However, when used with snapshots from an MD simulation, this same method outperforms all other methods selected in this study. This highlights the need for conformational sampling for reliable and efficient predictions of binding affinity for some systems. In our previous study, we combined coarse-grained forcefield with umbrella sampling to calculate ΔΔ*G* values for eight mutations of 3hfm Ab complex (one of the test systems in this study) and obtained better predictions than FoldX and MD+FoldX [56]. This study further emphasizes the need for better conformational strategies for some systems.

Statistical measures used to analyze performance are listed and defined in Table 3. For Ab, BPX+FoldX, MD+FoldX, and DCOMPLEX have the highest *r* values of the methods in our study (see Figure 4). MD+FoldX appears to be the most accurate method for Ab complexes. BindProfX, FoldX, JayZ, EasyE, and iSEE have low *r* and *ρ_c_* indicating that affinities estimated using these methods do not correlate well with experimental ΔΔ*G* values using a linear transformation. Also, the τ and *ρ* were lower compared to MD+FoldX, indicating these methods do poorly at calculating a rank order that matches experimental data.

The ROC curves allow us to determine each method’s ability to classify mutations as either destabilizing or neutral/stabilizing (Figures 3 and 5). For non-Ab complexes, JayZ (0.84 AUC) and EasyE (0.83 AUC) have the best true positive rate followed by DCOMPLEX (0.82 AUC). For Ab complexes, JayZ (0.97 AUC) and EasyE (0.98 AUC) have better true positive rates than MD+FoldX, the method with the highest *r* value. If classification of destabilizing vs stabilizing is the primary need, then JayZ or EasyE are both recommended over the other methods tested here due to their high accuracy and fast runtime.

While accuracy is generally the main reason for choosing a particular method, computational efficiency is also an important consideration, especially when predicting the effects of a large number of mutations. Here, we discuss the performance of each method in terms of its trade-off between speed and accuracy for predicting ΔΔ*G* values. For all single mutations and our non-Ab subset, EasyE and JayZ performed well; JayZ is the faster method of the two with EasyE at a similar speed to FoldX. DCOMPLEX is more accurate than FoldX for all single mutations and has similar accuracy as FoldX for non-Ab mutations, but at much lower cost. MD+FoldX has similar accuracy to DCOMPLEX for all single mutations and has similar accuracy to FoldX in non-Ab mutations but is by far the most computationally expensive method we tested. Although a synergistic combination of BPX+FoldX implements several structural and physico-chemical interaction terms in its algorithm, computation time was longer than all but MD+FoldX without a concomitant improvement in *r*. We note that this method is perhaps the most accessible of those tested, due to the easy-to-use online server interface and accuracy that is similar to FoldX for most systems. BindProfX utilizes the same scoring profile as BPX+FoldX without the FoldX calculations. In this case, accuracy decreased while calculation speed remained similar to BPX+FoldX. iSEE, the least correlating method, employs the widest variety of information to obtain relative binding affinity predictions and is the fastest of all methods (not including the non-trivial preparation time). For Ab complexes, MD+FoldX, the slowest of all the methods, had the highest accuracy, followed by DCOMPLEX. iSEE is again the fastest of all methods but also the least accurate. BindProfX utilizes several pre-calculated physico-chemical structural data in its scoring function while, JayZ and EasyE each layer an additional predictive calculating feature on top of Rosetta’s backbone sampling, adding complexity to the predictive algorithms. However, all three have similar *r* yet they do not achieve the accuracy of MD+FoldX. Overall, for non-Ab complexes, EasyE and JayZ appear to have the best balance between speed and accuracy of the methods we tested. For Ab complexes, DCOMPLEX appears to have the best balance.

We have demonstrated that all the tested methods have specific strengths and weaknesses; some perform better in specific contexts (Table 4), and some have longer runtimes to obtain similar predictive power to comparably faster methods. This highlights the complexity of the physico-chemical properties and structural features that drive, and limit, these predictive models. Our results can be used to make informed decisions for methods that may be preferable for a particular study or system. Table 4 suggests that if the goal is to estimate only the order of magnitude or sign of relative binding affinities, then the preferred method will likely be very different than if the goal is to obtain the best possible accuracy for antibody-antigen systems. To improve accessibility, we have generated an in-house Python script (provided in the supplement with the full dataset used in this work) that can be used to parse any of the parameters used in this study and provide tailored information. This information in combination with the runtime and other details provided in this study can be used to inform users on methods that can provide the best accuracy and efficiency for a given protein-protein complex type, set of physico-chemical features or structural parameters, and set of mutations. Additionally, the script can be extended to other methods and feature-sets, potentially elucidating specific problems or areas of improvement to existing and future methods.

## Conclusions

In this study, we have assessed the accuracy and efficiency of eight computational methods on predicting binding affinity changes due to single amino acid mutations. Methods were tested on 16 different protein complexes: eight antigen-antibody (Ab) and eight non-antigen-antibody (Non-Ab) complexes. While some methods perform consistently better than others, how well each performs depends on the physico-chemical and structural components of each complex. EasyE was the most accurate for non-Ab complexes, and MD+FoldX was most accurate for Ab complexes. JayZ and EasyE were better able to distinguish between destabilizing (ΔΔ*G* > 0.5 kcal/mol) and stabilizing (ΔΔ*G* < −0.5 kcal/mol) as compared to any other method. Future work could include more systems or different methods, including those that are solely web server-based in order to expand and better refine our conclusions on their predictive capability.

## Supporting information captions

**S1 File. A word document with detailed protocols for calculating ΔΔ*G* values using each of the eight methods used in this study**.

**S2 File. An in-house Python script that can be used to parse any of the parameters used in this study and provide tailored information**.

**S3 File. A CSV file with full dataset used in this work and predicted ΔΔ*G* values for each mutation using eight methods**.

**S1 Figure. Performance of each evaluated method for Ab and non-Ab complexes in predicting true ΔΔ*G* values (*ρ*_c_), linearly correlated ΔΔ*G* values (*r*), and rank order (*ρ* and τ) for a select subset of mutations that occur in beta sheet**. The error for each method is reported under the correlation points.

**S2 Figure. Performance of each evaluated method for Ab and non-Ab complexes in predicting true ΔΔ*G* values (*ρ*_c_), linearly correlated ΔΔ*G* values (*r*), and rank order (*ρ* and τ) for a select subset of mutations that occur in alpha helix**. The error for each method is reported under the correlation points.

**S3 Figure. Performance of each evaluated method for Ab and non-Ab complexes in predicting true ΔΔ*G* values (*ρ*_c_), linearly correlated ΔΔ*G* values (*r*), and rank order (*ρ* and τ) for a select subset of mutations with wild type amino acids that are either glycine or proline**. The error for each method is reported under the correlation points.

**S4 Figure. Performance of each evaluated method for Ab and non-Ab complexes in predicting true ΔΔ*G* values (*ρ*_c_), linearly correlated ΔΔ*G* values (*r*), and rank order (*ρ* and τ) for a select subset of mutations with wild type amino acids that are neither glycine nor proline**. The error for each method is reported under the correlation points.

